## Supplement for "Uncovering the mechanism for aggregation in repeat expanded RNA reveals a reentrant transition"

### Supplementary Information

### S1 Calculating the multiplicity factor $g$

In this section we describe the calculation of $g(n, m, N_b)$, the number of distinct possible combinations of $N_b$ bonds that can be formed with $m$ strands, each with $n$ stickers. We first derive $g(n, 1, N_b)$, i.e. the monomer case. We consider both the case in which adjacent stickers can bind to one another, and the case in which they cannot. We then consider multimers, deriving both exact and approximate expressions for $g(n, m, N_b)$. We also explain the origin of the $1/m$ term in $g(n, m, N_b)$, and its connection to symmetry factors.

#### S1.1 The calculation of $g$ for monomers

##### S1.1.1 Allowing neighbor binding

The question we pose here is as follows: given $n$ binding sites, each of which can be bound to at most one other binding site, how many non-pseudoknotted ways are there of constructing $N_b$ bonds between the sites?

There are $\binom{n}{2N_b}$ ways of choosing the $2N_b$ bound sites out of the $n$ possibilities. Then, there are $C_{N_b}$ distinct ways of constructing $N_b$ non-pseudoknotted bonds between $2N_b$ sites, where $C_i$ is the $i^{\text{th}}$ Catalan number, defined as

$$C_{N_b} = \frac{1}{N_b + 1}\binom{2N_b}{N_b}. \tag{S1}$$

Combining these expressions, we find that

$$g_a(n, 1, N_b) = \frac{n!}{(n - 2N_b)!\,(N_b + 1)!\,N_b!} \tag{S2}$$

where the subscript $a$ refers to the fact that the expression is the result of allowing neighbor binding.

##### S1.1.2 Disallowing neighbor binding

If the linker is too short to allow binding between adjacent stickers, the resulting expression is more complicated. To our knowledge, this case has not been explored previously, and so we derive it here.

We start with the result allowing neighbor binding, $g_a$ derived in the previous section. We find $g_d$ ("d" for "disallowing" neighbor binding) by taking $g_a$ and subtracting out the number of structures with at least one pair of neighboring stickers bound.

There are $n-1$ possible neighbor bonds, and for each, there are $g_a(n-2, 1, N_b-1)$ ways to arrange the remaining $N_b - 1$ bonds. There thus appear to be $(n-1)g_a(n-2, 1, N_b-1)$ ways to make $N_b$ bonds with $n$ stickers, including at least one neighbor bond. Our answer appears to be given by $g_a(n, 1, N_b) - (n-1)g_a(n-2, 1, N_b-1)$.

However, there is an error in that calculation, since some structures have two neighbor bonds! These structures were counted twice: once when fixing the first neighbor bond, and once when fixing the second. We therefore need to add back in the number of structures that have at least two neighbor bonds. Using a similar reasoning to previously, we get that the number of such structures appears to be $\frac{1}{2}(n-2)(n-3)g_a(n-4, 1, N_b-2)$. (We derive this result more fully below, but it perhaps makes intuitive sense: the division by two corrects for the fact that it doesn't matter in what order you determine the two bonds). Our answer thus appears to be

$$g_a(n, 1, N_b) - (n-1)g_a(n-2, 1, N_b-1) + \frac{(n-2)(n-3)}{2}g_a(n-4, 1, N_b-2). \tag{S3}$$

You may already see where this is going. That calculation itself had a similar error to the first: we now overcounted the number of structures with at least 3 neighboring bonds. Continuing this procedure further, we find that

$$g_d(n, 1, N_b) = g_a(n, 1, N_b) + \sum_{i=1}^{N_b} (-1)^i \binom{n-i}{i} g_a(n-2i, 1, N_b-i). \tag{S4}$$

The result of this sum is given by

$$g_d(n, 1, N_b) = \frac{(n-N_b)!\,(n-N_b-1)!}{(n-2N_b)!\,(n-2N_b-1)!\,(N_b+1)!\,N_b!} \tag{S5}$$

This result is exact and was validated by comparing to explicit computational enumeration.

To derive the factor of $\binom{n-i}{i}$, consider for example the case of $i = 2$: how many ways are there to specify two neighbor bonds? One of the two neighbor bonds comes first. If that bond is between the first and second stickers, there are $n-3$ ways to place the remaining neighbor bond. If that bond is between the second and third stickers, there are $n-4$ ways to place the second neighbor bond. Continuing on, there is only one place to place the remaining neighbor bond if the first bond is between the $(n-3)^{\text{rd}}$ and $(n-2)^{\text{nd}}$ stickers. Thus, the total number of ways to place the two bonds is

$$\sum_{j=1}^{n-3}(n-2-j) = \frac{(n-2)(n-3)}{2} = \binom{n-2}{2}. \tag{S6}$$

The general expression of $\binom{n-i}{i}$ can be derived along similar lines.

#### S1.2 The calculation of $g$ for multimers

##### S1.2.1 Allowing neighbor binding

If neighbors are allowed to bind, then the number of ways of constructing bonds between strands in a multimer is the same as the corresponding number for a monomer with the same total number of stickers. The result is therefore $g_a(n, m, N_b) = g_a(nm, 1, N_b)$.

One caveat here is that some of the bond combinations do not actually lead to a connected multimer. For example, $g_a(n, m, m-2)$ should always be zero – there is no way to make a multimer comprised of $m$ strands with $m-2$ bonds – but $g_a(nm, 1, m-2)$ need not be zero. Correcting the expression for $g$ to compute only connected multimers is the subject of a later section, but for the most part, this correction is negligible in the regimes we consider in this work.

Another caveat – of considering *distinct* structures, is also considered in a later section, and leads to a correction of $1/m$ in the expression above.

##### S1.2.2 Disallowing neighbor binding

If neighbors are not allowed to bind, then $g_d(nm, 1, N_b)$ slightly underestimates $g_d(n, m, N_b)$, even ignoring the two caveats mentioned above. This is because the last sticker of one strand is actually allowed to bind to the first sticker of the next, though this binding would be disallowed in a monomer. If we consider $g_d(nm + (m-1), 1, N_b)$, that is a slight *over*estimate: although it corrects for the previous issue, the new "phantom" stickers are allowed to bind in the monomer approximation, although they are not for the real multimer. An intermediate estimate, of $g_d(nm+\alpha(m-1), 1, N_b)$ appears to provide a reasonably good approximation for the true value of $g_d$ when $\alpha \approx 0.42$.

In the remainder of this section, we will derive an exact expression for $g_d(n, m, N_b)$. This derivation proceeds along similar lines to the derivation of the monomer case.

There are $g_a(n, m, N_b) = g_a(nm, 1, N_b)$ ways of making $N_b$ non-pseudoknotted bonds with $nm$ stickers, including neighbor pairings. There are $m(n-1)$ ways to fix one neighbor bond, and $g_a(nm-2, 1, N_b-1)$ ways to arrange the remaining bonds given one fixed bond. Therefore, it would appear that the result is $g_a(nm, 1, N_b) - m(n-1)g_a(nm-2, 1, N_b-1)$. However, some of the ways of rearranging the remaining bonds *themselves* have a neighbor bond, and so we counted those structures twice: once when fixing the first neighbor bond, and once when fixing the second. Following this through, as in the monomer case, we have that

$$g_d(n, m, N_b) = g_a(nm, 1, N_b) + \sum_{i=1}^{N_b} (-1)^i \, t(n, m, i) g_a(n-2i, 1, N_b-i) \tag{S7}$$

where $t(n, m, i)$ is the number of ways to fix $i$ neighbor bonds, given $m$ strands, each of length $n$. We found previously that $t(n, 1, i) = \binom{n-i}{i}$. For general $m$,

$$t(n,m,i) = \binom{m}{1} t(n,1,i) + \binom{m}{2} \sum_{j=1}^{i-1} t(n,1,j)t(n,1,i-j) + \binom{m}{3} \sum_{j=1}^{i-2} \sum_{k=1}^{i-2} t(n,1,j)t(n,1,k)t(n,1,i-j-k) + ... \quad \text{(S8)}$$

since there are $t(n,1,i)$ ways to fix $i$ neighbor bonds on a single strand and $\binom{m}{1}$ ways to pick a single strand out of $m$ strands; there are $t(n,1,j)$ ways to fix $j$ neighbor bonds on a single strand, $t(n,1,i-j)$ ways to fix the remaining bonds on another strand, and $\binom{m}{2}$ ways to choose the two strands; and so on.

While the sum in $t(n,m,i)$ can be written more succinctly in terms of generating functions, we are not aware of any simple closed-form formula for $g_d(n,m,N_b)$. We therefore rely on the heuristic described previously for our analytical calculations.

##### S1.2.3 A note about symmetry factors and $g$

In this work, we have used $g(n,m,N_b)$ to describe the number of *distinct* possible combinations of $N_b$ bonds that can be formed with $m$ strands, each with $n$ stickers. It is this italicized word *distinct* that leads to the factor of $1/m$ in $g(n,m,N_b)$ that was missing in the previous sections – it ensures identical structures are counted only once. However, this factor leads to results that are apparently non-sensical: it is possible for $g$ to result in a fractional output. As we will explain in this section, this is not an error, and in fact accounts for the symmetry-factor-based entropic cost of forming symmetric structures.

We begin by considering the expression for $g_a$ derived previously (identical arguments can be made for $g_d$). $g_a(nm,1,N_b)$ describes the number of – not necessarily distinct – possible combinations of $N_b$ bonds that can be formed with $m$ strands, each with $n$ stickers, treating an $m$-mer the same as a monomer $m$ times longer.

$g_a$ overcounts multimers comprised of $m$ strands by a factor of $m$. To demonstrate this, let us a consider one particular structure, for example, the structure shown in Fig. S1a. The panel shows a particular trimer structure of $n = 3$ CAG repeat strands. In panel b, we show that this trimer can be depicted in six (or 3!) different ways in the enumeration performed by $g_a$, corresponding to different permutations of the strands. However, three of these (shown on the right) have intersecting arcs and therefore are not considered as part of $g_a$'s enumeration. For 4-mers, there are 24 ways to enumerate each structure, but only 4 of these are considered within $g_a$ as the rest appear as pseudoknots. Thus, in order to count the contribution of each structure to the partition function only once, we need to divide the contribution of each $m$-mer structure to $Z$ by $m$. This corresponds to the number of cyclic permutations of the strands [1].

Some structures though are only enumerated once in $g_a$. In fact, structures that contain an R-fold symmetry are enumerated $m/R$ times. However, these should should indeed be counted with a partition function penalty of $1/R$ (or a free energy penalty of $k_BT\log(R)$) [1]. This symmetry factor correction arises from the entropic difference between structures with and without these symmetries [2]. An example of such a structure is shown in panels c-d of Fig. S1. The structure shown in panel c has 3-fold symmetry, and has only one possible depiction (panel d). Its free energy is thus effectively given by $3\Delta F_b - T\Delta S_{\text{loop}}(6) + 3\Delta G_{\text{assoc}} + k_BT\log(3)$. This latter term is effectively added by dividing its contribution to the partition function by 3 (the symmetry number). Thus, by dividing the contribution of each $m$-mer structure to $Z$ by $m$ we simultaneously correct for the overcounting performed by our enumeration procedure and account for the entropic penalty of symmetric structures.

##### S1.2.4 An exact calculation of $g$ for multimers

We will describe the exact calculation for $g(n,m,N_b)$ here. For most purposes, the exact calculation of $g$ is overkill: $g(n,m,N_b)$ is well approximated by $g(nm,1,N_b)/m$ (or a slight modification thereof when disallowing neighbor stickers from binding). However, in certain regimes – in particular for very small $n$ or very positive values of $F$ – this approximation is no longer valid. Moreover, we need to calculate $g$ exactly if we are to make the claim that the approximation we use is appropriate.

The issue we address in this section is that of *connected* multimers. To give an example (neglecting the factor of $1/m$ for the moment): $g_a(2,2,2)$ should be equal to 1; there is one way to make a dimer with two bonds, given two

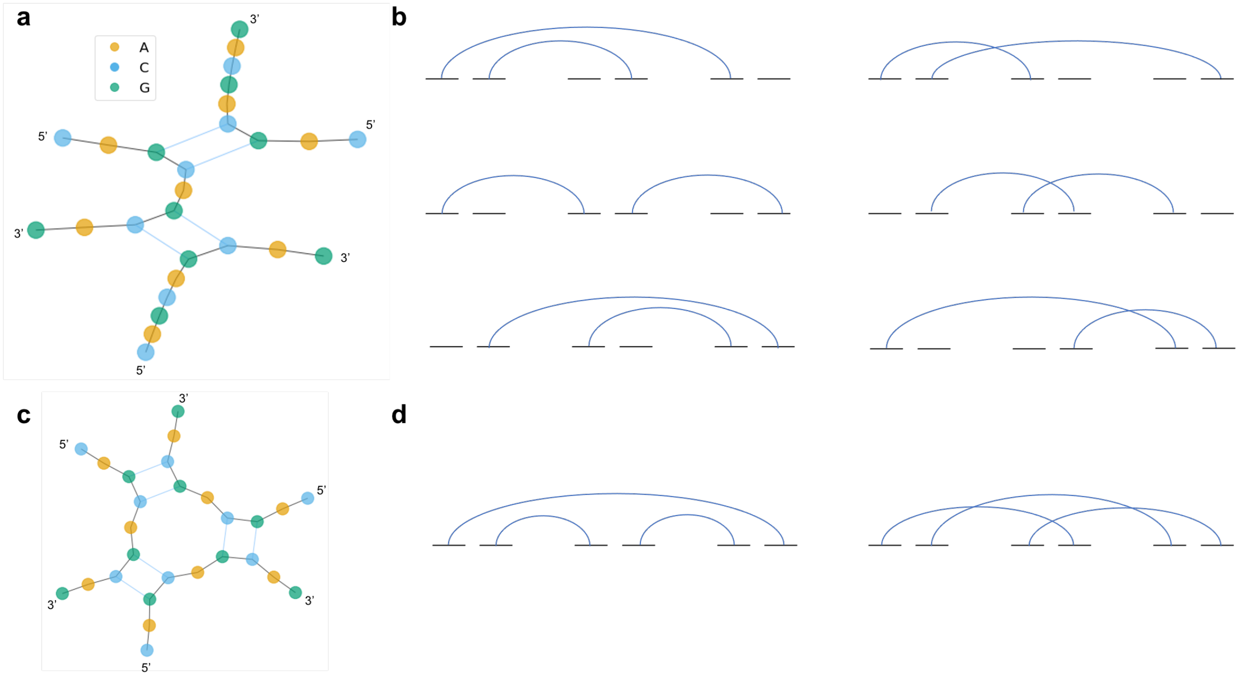

Figure S1: **Repeat RNA enumeration multiplicity**. **a:** A trimer of $n = 3$ CAG repeat strands as depicted by LandscapeFold is shown. **b:** Structures can be depicted by treating each sticker as a site (black lines) that can be connected to at most one other site (blue arcs show connections). The structure from panel a can be depicted in six ways corresponding to permutations of the strands. Three of the ways (right column) appear pseudoknotted (they have intersecting arcs) though the structure contains no pseudoknots. **c:** Another trimer of $n = 3$ CAG repeats is shown. **d:** This trimer can only be depicted in two ways in the abstraction, one of which appears pseudoknotted. See main text for discussion.

strands each with two stickers. (Accounting for the symmetry factor, the structure has a two-fold symmetry, so that $g_a/m = 1/2$). However, $g_a(4, 1, 2) = 2$ is twice as large, since it also considers the structure which has each sticker bound to its neighbor on the same strand (a structure that is also two-fold symmetric). This second structure is not a dimer in truth – it is actually two monomers. This problem gets worse with larger $m$ (for very small $n$). For example, $g_a(2, 3, 3)$ should be equal to 1: there is only one way to make a trimer with three bonds between three strands of $n = 2$ (and in fact, this trimer is shown in panels c and d of Fig. S1). However, $g(6, 1, 3) = 5$ since it also considers the structure comprised of 3 monomers, as well as the three permutations of the same structure comprised of one dimer and one monomer.

Typically, corrections of this sort are negligible, since the vast majority of possible structures are connected. However, in the regime that very few stickers are typically bound, or that all stickers are bound with a very small value of $n$ (such as $n = 2$), this is no longer the case.

An exact calculation of $g(n, m, N_b)$ therefore subtracts out the contribution of disconnected structures. We thus need to calculate all integer partitions of $m$, and for each, the number of ways to connect $m$ strands (excluding connections that appear as pseudoknots when drawing the strands in a line, as those are already not counted). For example, for $m = 4$, there is 1 way to connect the strands as 4 monomers, 6 ways to connect them as 2 monomers and a dimer, 4 ways to connect them as a monomer and a trimer, and 2 ways to connect them as 2 dimers. We then need to enumerate, for each of these combinations, the different ways of splitting up the $N_b$ bonds among the different sub-structures.

Thus, in order to calculate the connected multiplicity factor $g_{\text{conn}}(n, 4, N_b)$, we calculate:

$$
\begin{aligned}
g_{\text{conn}}(n,4,N_b) = g_{a/d}(n,4,N_b) - \Bigg[ & \sum_{i=0}^{N_b-j-k-l} \sum_{j=0}^{N_b-k-l} \sum_{k=0}^{N_b-l} \sum_{l=0}^{N_b} g(n,1,i)g(n,1,j)g(n,1,k)g(n,1,l) + \\
& 6 \sum_{i=0}^{N_b-j-k} \sum_{j=0}^{N_b-k} \sum_{k=0}^{N_b} g(n,1,i)g(n,1,j)g_{\text{conn}}(n,2,k) + \\
& 4 \sum_{i=0}^{N_b-j} \sum_{j=0}^{N_b} g(n,1,i)g_{\text{conn}}(n,3,j) + \\
& 2 \sum_{i=0}^{N_b-j} \sum_{j=0}^{N_b} g_{\text{conn}}(n,2,i)g_{\text{conn}}(n,2,j) \Bigg].
\end{aligned}
\tag{S9}
$$

Here, $g_{a/d}$ is the unconnected multiplicity as defined in Sections S1.2.1 and S1.2.2 for the cases of allowing and disallowing neighbor binding, respectively (the former case simply yields $g_a(4n, 1, N_b)$; the latter is more complicated as discussed in the referenced section). This unconnected multiplicity is corrected by subtracting the contribution of 4 monomers that have among them $N_b$ total bonds, the contribution of 2 monomers and 1 dimer, and so on.

Finally, to get the true multiplicity factor, $g_{\text{conn}}(n, m, N_b)$ needs to be divided by $m$ to account for symmetries as discussed in Section S1.2.3.

#### S2 Concentration units and their connection to $\Delta F$

Throughout the text, we have treated concentrations as dimensionless. In other words, the concentrations we use are normalized by some reference concentration $\rho$. We have not defined $\rho$ explicitly, because its value does not affect the physical observables we consider here. As explained in Ref. [1], any reference concentration used also enters into the definition of $\Delta G_{\text{assoc}}$, as

$$\Delta G_{\text{assoc}} = \Delta G^{\text{pub}}_{\text{assoc}} - \frac{1}{\beta}\log\left(\frac{\rho}{1\ \text{mol/Liter}}\right) \tag{S10}$$

where $\Delta G^{\text{pub}}_{\text{assoc}}$ is the published value for the free energy cost of dimerization. This value is $4.09$ kcal/mol for the free energy cost of RNA-RNA association [3, 4, 5]; $1.96$ for the free energy cost of DNA-DNA association [6]; and $3.1$ for the free energy cost of RNA-DNA association [7, 8].

The effect is that factors of $\rho$ cancel out in the relevant equations. Namely,

$$\begin{aligned}
\frac{c_m/\rho}{(c_1/\rho)^m} &= \frac{Z_m}{Z_1^m} \\
\frac{c_m}{c_1^m}\rho^{m-1} &= \frac{Z'_m e^{-\beta(m-1)\Delta G_{\text{assoc}}}}{Z_1'^m} \\
\frac{c_m}{c_1^m}\rho^{m-1} &= \frac{Z'_m e^{-\beta(m-1)\Delta G^{\text{pub}}_{\text{assoc}}}}{Z_1^m}\left(\frac{\rho}{1\ M}\right)^{m-1}
\end{aligned} \tag{S11}$$

where $Z'_m$ is $Z_m$ not including the free energy cost of multimerization.

In fact, the natural units with which to measure concentration are $c/(e^{\beta\Delta G^{\text{pub}}_{\text{assoc}}} \times 1\ M)$, since the above equation could be further simplified to

$$\frac{c_m/e^{\beta\Delta G^{\text{pub}}_{\text{assoc}}}}{(c_1/e^{\beta\Delta G^{\text{pub}}_{\text{assoc}}})^m} = \frac{Z'_m}{Z_1^m} \tag{S12}$$

where $c$ is now made dimensionless by measuring it in units of $M$, and the right-hand side is now independent of $G^{\text{pub}}_{\text{assoc}}$.

We chose to keep $\Delta G_{\text{assoc}}$ explicitly in the equations, in order to emphasize that changes to its value (for example, by changing ionic conditions) can affect aggregation properties. However, you will notice that both concentrations and $\Delta G^{\text{pub}}_{\text{assoc}}$ always appear in the combination $c/e^{\beta\Delta G^{\text{pub}}_{\text{assoc}}}$. Since we change units early on to $\Delta F \equiv \Delta G^{\text{pub}}_{\text{assoc}} - T\Delta S_{\text{loop}}$, that combination becomes $c/e^{\beta\Delta F}$.

### S3 Calculating the partition function analytically

The partition function is given by

$$Z_m = \sum_{N_b} g(n, m, N_b) e^{-\beta F N_b} e^{-\beta(m-1)\Delta F}. \tag{S13}$$

In principle, we can of course compute this sum. In practice however, such a computation is too computationally intensive for reasonable purposes. We therefore need to approximate the sum. We do so by a saddlepoint approximation, approximating the sum as being dominated by a particular term. We allow for a maximum of a second order correction in our approximations in order to balance computational feasibility and accuracy of the model. Indeed, these approximations appear to be entirely sufficient to describe the system with a high degree of accuracy.

In this section, we will describe the different regimes of this saddlepoint approximation.

#### S3.1 Allowing neighbor bonds

In this case, for $n \gg 1$, a good approximation for $g$ is

$$g(n, m, N_b) = \frac{(nm)!}{m\ (nm - 2N_b)!\ (N_b + 1)!\ N_b!} \tag{S14}$$

(See section S1.2.4 for a description of the calculation for very small values of $n$). In order to proceed, we need an estimate of which term in the sum is dominant.

##### S3.1.1 (Almost) all stickers are typically bound

The maximum possible number of stickers bound is

$$N_b^{\text{max}} = \text{floor}\left(\frac{nm}{2}\right). \tag{S15}$$

Finding an approximation for $g$ therefore depends on whether $nm$ is even or odd.

**Even** $nm$
In this case, $N_b^{\text{max}} = nm/2$. We then have

$$g(n, m, N_b^{\text{max}}) = \frac{(nm)!}{m\ \left(\frac{nm}{2} + 1\right)\ \left[\left(\frac{nm}{2}\right)!\right]^2} \tag{S16}$$

where we have written the factorial in a way that will make the final result cleaner. We approximate the factorial with Stirling's approximation, $x! \approx \sqrt{2\pi x}(x/e)^x$, yielding

$$g(n, m, N_b^{\text{max}}) \approx \frac{2^{nm}}{(nm + 2)\sqrt{\frac{n\pi}{8}} m^{3/2}} \tag{S17}$$

**Odd** $nm$
In this case, $N_b^{\text{max}} = (nm - 1)/2$. We then have

$$g(n, m, N_b^{\text{max}}) = \frac{nm(nm - 1)!}{m\ \left(\frac{nm+1}{2}\right) \left[\left(\frac{nm-1}{2}\right)!\right]^2} \tag{S18}$$

where, again, the factorials have been broken up to make the final expression cleaner. Using Stirling's approximation, we get

$$g(n, m, N_b^{\text{max}}) \approx \frac{2^{nm-1}}{\frac{nm+1}{n}\sqrt{\frac{(nm-1)\pi}{8}}} \tag{S19}$$

We can then repeat this procedure to get the next order correction, meaning the term corresponding to $N_b^{\text{max}} - 1$ bonds. All combined, we find that

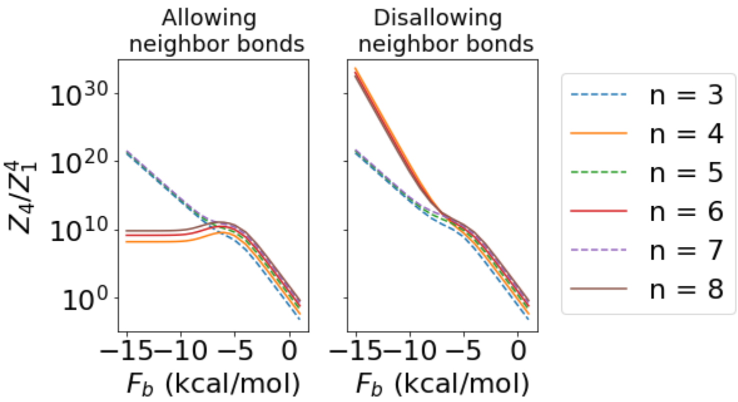

Figure S2: **Partition functions as a function of** $F_b$. $Z_4/Z_1^4$ is plotted as a function of $F_b$ using the exact computationally-enumerated partition functions. The surprising behavior predicted by the analytical model and discussed in the main text can be seen clearly, namely that at the interface between the strong and intermediate binding regimes for even $n$, the partition function ratio is larger than it is in the very strong binding regime. This behavior is seen regardless of the value of $m$ chosen (here, $m = 4$).

$$Z_m \approx \begin{cases} e^{\left(\log(2)-\frac{\beta F}{2}\right)nm-\beta(m-1)\Delta F}\,\frac{1}{m}\left(\frac{1}{nm+2}\sqrt{\frac{8}{nm\pi}}+\sqrt{\frac{nm}{8\pi}}e^{\beta F}+\mathcal{O}\left(e^{2\beta F}\right)\right) & \text{if } nm \text{ is even} \\ e^{\left(\log(2)-\frac{\beta F}{2}\right)(nm-1)-\beta(m-1)\Delta F}\,\frac{1}{m}\left(\frac{nm}{nm+1}\sqrt{\frac{8}{(nm-1)\pi}}+\frac{nm}{3}\sqrt{\frac{nm-1}{8\pi}}e^{\beta F}+\mathcal{O}\left(e^{2\beta F}\right)\right) & \text{otherwise} \end{cases} \quad \text{(S20)}$$

in this regime.

For very negative $\beta F$, the first term is dominant. For even values of $n$, that implies that $Z_m/Z_1^m$ is independent of $F$ for very negative values. Furthermore, for intermediate values of $\beta F$ and even $n$, $Z_m/Z_1^m$ is actually larger than for very negative $\beta F$. This behavior is at the heart of the reentrant transition described in the main text, and can be seen in the exact computationally-enumerated partition function in Fig. S2

##### S3.1.2 Calculating $N_b^\star$

For the regime in which some stickers are typically bound (next section) we need to calculate the dominant term of $Z_m$, corresponding to $N_b^\star$. This corresponds to the value of $N_b$ such that

$$\frac{\partial g(n,m,N_b)e^{-\beta F N_b}}{\partial N_b} = \frac{\partial}{\partial N_b}\left[\frac{(nm)!\, e^{-\beta F N_b}}{m\,(nm-2N_b)!\,(N_b+1)!\,N_b!}\right] = 0. \quad \text{(S21)}$$

Calculating the derivative and simplifying, we arrive at

$$\frac{1}{N_b^\star+1} + \beta F + 2\psi^{(0)}(N_b^\star+1) = 2\psi^{(0)}(nm-2N_b^\star+1) \quad \text{(S22)}$$

where $\psi^{(m)}(z)$ is the polygamma function of order $m$, defined in terms of the gamma function as

$$\psi^{(m)}(z) = \frac{d^{m+1}}{dz^{m+1}}\log\Gamma(z). \quad \text{(S23)}$$

For large values of $z$, $\psi^{(0)}(z+1)$ is approximately given by $\log(z)$. This approximation directly applies in the intermediate binding regime: the argument of the left-hand-side polygamma function is large when not in the weak binding regime, and the argument of the right-hand-side function is large when not in the strong binding regime. Assuming the $N_b^\star \gg 1$, we also neglect the first term, arriving at

$$\beta F + 2\log(N_b^\star) = 2\log(nm-2N_b^\star). \quad \text{(S24)}$$

Solving this equation, we have

$$N_b^\star = \frac{nm}{2 + e^{\beta F/2}} \tag{S25}$$

##### S3.1.3 Some stickers are typically bound

As derived above, we find

$$N_b^\star \approx \frac{nm}{2 + e^{\beta F/2}}. \tag{S26}$$

When $e^{\beta F/2} \ll 1$, the previous regime applies, and the discreteness of the number of stickers matters. In the regime with which we are concerned here, the fact that $N_b^\star$ is not an integer is insignificant, and in fact, as we will see, the sum can be well approximated as an integral.

This regime applies when each of the factorial terms in (S14) is greater than unity. We therefore approximate the factorial with Stirling's approximation, $x! \approx \sqrt{2\pi x}(x/e)^x$, yielding

$$g(n, m, N_b) \approx \frac{1}{2\pi m N_b(N_b+1)} \left(\frac{nm}{nm - 2N_b}\right)^{nm+\frac{1}{2}} \left(\frac{nm - 2N_b}{nm}\right)^{2N_b} \tag{S27}$$

Plugging in $N_b^\star$ for $N_b$, we have

$$g(n, m, N_b) \approx \frac{1}{2\pi m N_b^\star(N_b^\star+1)} \left(1 + 2e^{-\beta F/2}\right)^{nm+\frac{1}{2}} \left(e^{\beta F/2}\right)^{2N_b^\star} \tag{S28}$$

Thus, the dominant term of $Z_m$, (which we call $Z_m^\star$) is approximately

$$Z_m^\star \approx \frac{1}{2\pi m N_b^\star(N_b^\star+1)} \left(1 + 2e^{-\beta F/2}\right)^{nm+\frac{1}{2}} e^{-\beta(m-1)\Delta F} \tag{S29}$$

which can be written more tellingly as

$$Z_m^\star \approx \frac{e^{\beta\Delta F}\left(1 + 2e^{-\beta F/2}\right)^{1/2}}{2\pi m N_b^\star(N_b^\star+1)} \left[\left(1 + 2e^{-\beta F/2}\right)^{n} e^{-\beta\Delta F}\right]^m \tag{S30}$$

Written this way, connections between this expression and that found in the previous regime are apparent. In particular, it is clear why $\log Z$ increases linearly with both $n$ and $m$, and the factor of $2^{nm}$ makes an appearance here as in the previous regime. However, understanding the behavior of $Z_m/Z_1^m$ involves understanding the behavior of the prefactor to the bracketed term.

We also consider the next-order correction to $Z_m$. This is found using the saddlepoint approximation. For an exponential integrand $e^{f(x)}$ that has a maximum at $x^\star$ (and therefore $f'(x^\star) = 0$), the following becomes a very good approximation since exponentials are so sharply peaked:

$$\int_{-\infty}^{\infty} e^{f(x)}dx \approx \int_{-\infty}^{\infty} e^{f(x^\star)+xf'(x^\star)+\frac{x^2}{2}f''(x^\star)}dx = e^{f(x^\star)}\sqrt{\frac{2\pi}{|f''(x^\star)|}}. \tag{S31}$$

For our purposes, $f(N_b) = \log\left(g(n, m, N_b)\right) - \beta F N_b - \beta(m-1)\Delta F$. What we have done so far – finding $Z_m^\star$ – is equivalent in this language to finding $e^{f(x^\star)}$. Since $|f''(N_b^\star)|$ is generally quite small ($\mathcal{O}(1)$ as a general rule, $< 6$ in all cases we examined), the error introduced by the approximation of the sum as an integral is negligible. In this case, the curvature term $f''(x^\star)$ leads to

$$Z_m \approx Z_m^\star \sqrt{\frac{2\pi}{4\psi^{(1)}(n - 2N_b^\star + 1) + 2\psi^{(1)}(N_b^\star + 1) - \frac{1}{\left(N_b^\star+1\right)^2}}}. \tag{S32}$$

We compare this analytical formula to the computational results for intermediate binding ($F_b = -6$ kcal/mol) in Fig. S3. Here we use the parameter $l_{\text{eff}}$ fit to data from Fig. 2 along with (9).

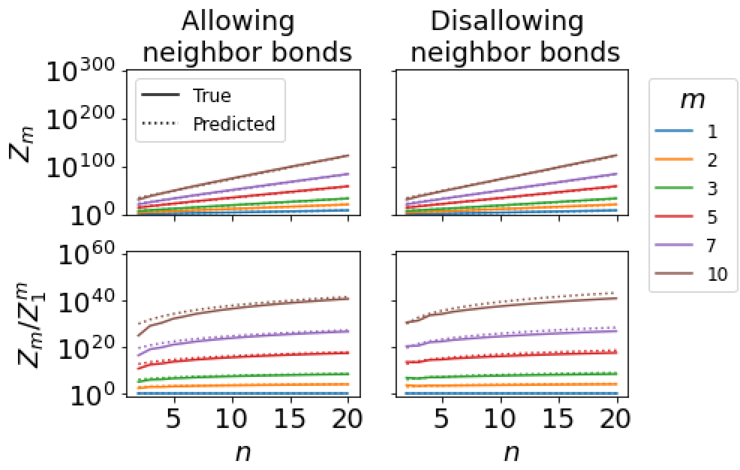

Figure S3: **Partition functions in the intermediate regime**. $Z_m$ and $Z_m/Z_1^m$ are plotted as in Fig. 2 with $F_b = -6$ kcal/mol. A single parameter for each column is fit to the results in Fig. 2 as described in the discussion of that figure, and extrapolated to $F_b = -6$ kcal/mol using (9). As predicted, there is no even/odd discrepancy for $n$ in this intermediate regime of binding.

###### S3.1.4 Very few stickers are typically bound

This is the regime in which the typical number of stickers in a multimer of size $m$ is equal to or only slightly larger than $m-1$. This is the most computationally difficult regime to explore, since in this regime, $g(n,m,N_b)$ is not well approximated by $g(nm,1,N_b)/m$. Instead, we need to employ the exact calculation of $g(n,m,N_b)$ described in Section S1.2.4. We consider only the first three terms of the sum in this regime, enumerating the contribution of $N_b = m-1$ up to $m-3$. We do not consider this regime further analytically, but do employ it in comparison to the computational predictions (e.g. Fig. 2B).

#### S3.2 Disallowing neighbor bonds

In this case, a good approximation for $g(n,m,N_b)$ is given by

$$g(n,m,N_b) \approx \frac{(q-N_b)!\,(q-N_b-1)!}{m\,(q-2N_b)!\,(q-2N_b-1)!\,(N_b+1)!\,N_b!} \tag{S33}$$

where $q = nm + \alpha(m-1)$, with $\alpha \approx 0.42$.

We will follow a similar procedure in this section as that taken in the previous section, but the calculations are slightly more cumbersome here because of the increased complexity of $g$.

###### S3.2.1 (Almost) all stickers are typically bound

In this regime, we find that we get a good approximation for $Z$ by considering the monomer case, and substituting in $q$ for $n$ after simplifying.

With neighbor bonds disallowed, the maximum possible number of stickers bound for monomers is

$$N_b^{\max} = \text{floor}\left(\frac{n-1}{2}\right). \tag{S34}$$

Finding an approximation for $g$ therefore again depends on whether $n$ is even or odd. The procedure here is analogous – with slightly more involved calculations – to the procedure for calculating $Z$ allowing for neighbor binding.

**Odd** $n$

The final term in the sum for $Z_1$ for odd $n$ is given by $N_b = \frac{n-1}{2}$. After simplifying the expression for $g$ in this case, we find that there is only one possible way to arrange this number of bonds in a monomer. The next term corresponds to a value of $N_b$ given by $\frac{n-3}{2}$. Combined, we find that in this regime,

$$Z_1 = e^{-\frac{\beta F}{2}(n-1)} \left(1 + \frac{(n+3)(n+1)^2(n-1)}{192} e^{\beta F}\right). \tag{S35}$$

**Even $n$**

The final term in the sum for $Z_1$ for even $n$ is given by $N_b = \frac{n}{2} - 1$. Simplifying the factorials and including the second-to-last term as well, we find that for even $n$ in this regime,

$$Z_1 = e^{-\frac{\beta F}{2}(n-2)} \left(\frac{n(n+2)}{8}\right) \left(1 + \frac{(n+4)(n+2)n(n-2)}{1152} e^{\beta F}\right). \tag{S36}$$

To convert from $Z_1$ to $Z_m$, we first multiply by $\frac{e^{-\beta(m-1)\Delta F}}{m}$. Second, we (heuristically) replace each instance of $n$ in these expressions by $q$. Finally, for even values of $nm$ with $m > 1$, multimers are described by this equation with the exception that they are able to form one further bond (such that all stickers are bonded). We can account for this by adding a term

$$\frac{e^{-\beta(m-1)\Delta F}}{m} e^{-\frac{\beta F}{2} q} \tag{S37}$$

to the partition function for even $n$. This term is primarily useful for $m = 2$, for which the multiplicity factor of one is accurate (for $m = 2$, there is indeed only one way to fulfill all bonds, when disallowing neighbor binding). For $m > 2$, the replacement of $n$ by $q$ partially accounts for the lack of a further multiplicity factor; furthermore, we ultimately find that multimers for $m > 2$ are fairly well approximated by the intermediate regime – considered in the following sections – for the free energies we consider.

##### S3.2.2 Calculating $N_b^\star$

For the regime in which some stickers are typically bound (next section) we need to calculate the dominant term of $Z_m$, corresponding to $N_b^\star$. This corresponds to the value of $N_b$ such that

$$\frac{\partial g(n,m,N_b)e^{-\beta F N_b}}{\partial N_b} = \frac{\partial}{\partial N_b}\left[\frac{((q-N_b)!)^2\ (q-2N_b)\ e^{-\beta F N_b}}{m\ (q-N_b)\ ((q-2N_b)!)^2\ (N_b+1)\ (N_b!)^2}\right] = 0. \tag{S38}$$

Calculating the derivative and simplifying, we arrive at

$$\frac{2}{q-2N_b^\star} + \frac{1}{N_b^\star+1} - \frac{1}{q-N_b^\star} + \beta F - 4\psi^{(0)}(q-2N_b^\star+1) + 2\psi^{(0)}(q-N_b^\star+1) + 2\psi^{(0)}(N_b^\star+1) = 0. \tag{S39}$$

Recognizing that in the intermediate regime, $q - 2N_b^\star > 1$, $N_b^\star + 1 > 1$, and $q - N_b^\star > 1$, we can make the approximation of neglecting the first three terms. We then make the same approximation as previously, treating $\psi^{(0)}(z+1) \approx \log(z)$. Solving the resulting equation and recognizing that $N_b^\star < \frac{q}{2}$ to remove the extraneous solution, we arrive at

$$N_b^\star \approx \frac{q}{2}\left(1 - \frac{e^{\beta F/4}}{\sqrt{4+e^{\beta F/2}}}\right). \tag{S40}$$

##### S3.2.3 Some stickers are typically bound

As derived above, we find that for monomers

$$N_b^\star \approx \frac{n}{2}\left(1 - \frac{e^{\beta F/4}}{\sqrt{4+e^{\beta F/2}}}\right). \tag{S41}$$

where, as previously, we substitute $q$ for $n$ when considering multimers.

The dominant term of $Z$ for monomers corresponds to

$$g \approx \frac{(n-2N_b^\star)\left[(n-N_b^\star)!\right]^2}{(n-N_b^\star)(N_b^\star+1)\left[(n-2N_b^\star)!\right]^2\left[(N_b^\star)!\right]^2}. \tag{S42}$$

Using Stirling's approximation, this expression becomes

$$g \approx \frac{1}{2\pi m N_b^\star(N_b^\star+1)}\left(\frac{n-N_b^\star}{n-2N_b^\star}\right)^{2(n-N_b^\star)}\left(\frac{n}{N_b^\star}-2\right)^{2N_b^\star}. \tag{S43}$$

As previously, we consider not only the dominant term of $Z$ but also the curvature term of the saddlepoint approximation. This term can be most clearly written in terms of the following function

$$h(x) = x^{-2} - 2\psi^{(1)}(x) \tag{S44}$$

as

$$\sqrt{\frac{2\pi}{-4h(n-2N_b^\star)+h(n-N_b^\star)-h(N_b^\star+1)}}. \tag{S45}$$

Putting this all together, in this regime, we approximate $Z_m$ as

$$Z_m \approx Z_m^\star\sqrt{\frac{2\pi}{-4h(q-2N_b^\star)+h(q-N_b^\star)-h(N_b^\star+1)}} \tag{S46}$$

where

$$\begin{aligned} Z_m^\star =& \frac{e^{-\beta(FN_b^\star+(m-1)\Delta F)}}{2\pi m N_b^\star(N_b^\star+1)}\left(\frac{q-N_b^\star}{q-2N_b^\star}\right)^{2(q-N_b^\star)}\left(\frac{q}{N_b^\star}-2\right)^{2N_b^\star} \\ N_b^\star =& \frac{q}{2}\left(1-\frac{e^{\beta F/4}}{\sqrt{4+e^{\beta F/2}}}\right). \end{aligned} \tag{S47}$$

### S4 Predicting the aggregation threshold

Once we have the partition functions, we can then compute the equilibrium concentration of each multimer. These are found by solving the following set of equations:

$$\begin{aligned} c_m &= \frac{Z_m}{Z_1^m} c_1^m \\ \sum_m m c_m &= c^{\text{tot}} \end{aligned} \tag{S48}$$

where the concentrations are made dimensionless by normalizing by a reference concentration (see Section S2) and $c^{\text{tot}}$ is the total concentration of strands added to solution.

Aggregation is predicted when $c_m$ grows with $m$. However, these equations have $c_m$ dependent on $c_1$. If $m$ has a finite maximum value $m_{\text{max}}$, we can simply solve the $m_{\text{max}}$ equations for $c_1$, plug that in, and observe the dependence on $m$ of $c_m$. Indeed, this is how we make predictions of $c_m$ in our computational model. There is also some finite value to $m_{\text{max}}$ in an experiment given by the total number of molecules in the solution. However, that number is far too large to treat *via* this approach, and is reasonably treated as infinite. How can we find the aggregation threshold allowing for arbitrarily large clusters?

In this section, we will demonstrate how this question can be addressed. We will first consider the case allowing neighbor binding, and then the case disallowing neighbor binding. For each, the simplest case to consider is that in which monomers have some of their stickers typically bound (i.e. the intermediate regime considered above), and that is where we will begin.

Throughout this section, we will consider only the dominant term of the sum for $Z_m$, without any higher-order corrections. Some accuracy is therefore compromised for the sake of computational feasibility, but errors are expected to be minor.

#### S4.1 General framework

We start with a simple example to demonstrate the framework of the calculation. Consider a partition function

$$Z_m = \kappa m^{-\lambda} \gamma^m. \tag{S49}$$

Partition functions for this system are often approximately in this form. The concentration of $m$-mers $c_m$ is given in terms of $c_1$ by

$$c_m = Z_m \left(\frac{c_1}{Z_1}\right)^m = \kappa m^{-\lambda} \left(\frac{c_1}{\kappa}\right)^m. \tag{S50}$$

The total concentration is given by

$$c^{\text{tot}} = \sum_m m c_m = \kappa \sum_m m^{-\lambda+1} \left(\frac{c_1}{\kappa}\right)^m. \tag{S51}$$

We now recognize that the right-hand-side has a maximum possible value it can reach for $c_1 \leq \kappa$, given by $\kappa\zeta(\lambda - 1)$ where $\zeta$ is the Riemann zeta function:

$$\zeta(s) = \sum_{n=1}^{\infty} \frac{1}{n^s}. \tag{S52}$$

if $c^{\text{tot}} > \kappa\zeta(\lambda - 1)$, no value of $c_1 \leq \kappa$ will satisfy (S51). In that case, $c_1/\kappa$ must be $> 1$, and thus $c_m$ increases with $m$ for large enough $m$. The aggregation threshold is thus defined by

$$c^{\text{tot}}_{\text{thresh}} = \kappa\zeta(\lambda - 1). \tag{S53}$$

For concentrations greater than $c^{\text{tot}}_{\text{thresh}}$, the system is expected to form aggregates; for smaller concentrations, $c_m$ exponentially decreases with $m$.

To be clear, for $c^{\text{tot}} > c^{\text{tot}}_{\text{thresh}}$ (i.e. $c_1 > \kappa$) the sum in (S51) will actually diverge, an outcome that is both unphysical and unreasonable mathematically given that the left hand side is finite. The solution is that we are neglecting here any excluded volume interactions which will contribute a term $e^{-vm^2}$ in the summand and ensure the sum does not actually diverge, and physically mean that we will still expect to form finite-sized clusters in the aggregation regime. However, since we expect $v$ to be small, the integrand will still increase with $m$ within a certain regime for $c_1/\kappa > p$, where $p$ is only fractionally larger than unity. We will continue to neglect these effects here, since they are not, for our purposes, instructive.

#### S4.2 Allowing neighbor bonds

##### S4.2.1 Monomers have some stickers typically bound

The number of stickers typically bound is approximately

$$N_s(m) \approx \frac{2nm}{2 + e^{\beta F/2}} \tag{S54}$$

Here we are treating values of $n$ and $\beta F$ such that $n - 2 > N_s(1) > 6$ (or so) such that the intermediate regime approximation applies for monomers.

A question arises: if monomers are in the intermediate regime, how about multimers of size $m$? Because $N_s(m)$ scales linearly with $m$, a constant fraction of sites are expected to be bound, regardless of $m$. Therefore, the intermediate regime will always apply for multimers, if it applies for monomers.

We found previously that the partition function in this regime is approximately given by

$$Z_m = \frac{e^{\beta\Delta F}\left(1 + 2e^{-\beta F/2}\right)^{1/2}}{2\pi m N_b^\star (N_b^\star + 1)} \left[\left(1 + 2e^{-\beta F/2}\right)^n e^{-\beta\Delta F}\right]^m. \tag{S55}$$

The bracketed term cancels exactly in the equation for $Z_m/Z_1^m$, yielding

$$\frac{Z_m}{Z_1^m} = \frac{1}{m^2}\left[\frac{e^{\beta\Delta F}\left(1 + 2e^{-\beta F/2}\right)^{1/2}\left(2 + e^{\beta F/2}\right)^2}{2\pi n\left(n + 2 + e^{\beta F/2}\right)}\right]^{1-m} \frac{n + 2 + e^{\beta F/2}}{nm + 2 + e^{\beta F/2}}. \tag{S56}$$

We can then combine equations (S48) with this to get

$$c^{\text{tot}} = \frac{e^{\beta\Delta F}\left(1 + 2e^{-\beta F/2}\right)^{1/2}\left(2 + e^{\beta F/2}\right)^2}{2\pi n} \sum_{m=1}^{\infty} \frac{1}{m\left(nm + 2 + e^{\beta F/2}\right)} \left[x(c_1)\right]^m \tag{S57}$$

where we have defined $x(c_1)$ to be

$$x(c_1) = \frac{2\pi n\left(n + 2 + e^{\beta F/2}\right) c_1}{e^{\beta\Delta F}\left(1 + 2e^{-\beta F/2}\right)^{1/2}\left(2 + e^{\beta F/2}\right)^2} \tag{S58}$$

We will discuss in the next section a way to use this equation as is, but for now, in order to proceed will will make the assumption that $nm \gg 2 + e^{\beta F/2}$. This is equivalent to the assumption that $N_b^\star \gg 1$, which is entirely reasonable in this regime. This allows us to write

$$c^{\text{tot}} = \sum_m m\, c_m \approx \frac{e^{\beta\Delta F}\left(1 + 2e^{-\beta F/2}\right)^{1/2}\left(2 + e^{\beta F/2}\right)^2}{2\pi n^2} \sum_{m=1}^{\infty} \frac{\left[x(c_1)\right]^m}{m^2} \tag{S59}$$

Since the prefactor to the sum is independent of $m$, the statement that $c_m$ increases with $m$ (our definition of aggregation) is equivalent to the statement that $x(c_1) > 1$. Once $x(c_1) > 1$, then for large enough $m$, the summand will be increasing with $m$.

We have now refined our problem of finding the aggregation threshold as finding the set of parameters for which $x(c_1) > 1$. However, we still don't have an estimate for $c_1$! How then can we find for what set of parameters $x(c_1) > 1$?

Let us rewrite (S59) in a more suggestive form:

$$\frac{2\pi n^2 c^{\text{tot}}}{e^{\beta\Delta F}\left(1 + 2e^{-\beta F/2}\right)^{1/2}\left(2 + e^{\beta F/2}\right)^2} = \sum_{m=1}^{\infty} \frac{\left[x(c_1)\right]^m}{m^2} \tag{S60}$$

The key comes from recognizing that for $x(c_1) \leq 1$, the right hand side has a maximum value it can reach, given by the Riemann zeta function $\zeta(2) = \pi^2/6$. If the left hand side is greater than that value, $x(c_1)$ *must be* $> 1$. (And again – the reason the sum nonetheless evaluates to a finite value is because of the excluded volume interactions we are omitting here). Since the left hand side is comprised only of terms we control directly – $c^{\text{tot}}$, $n$, $\beta F$, and $\beta\Delta F$ – we have found our condition for aggregation in this regime:

$$\frac{n^2 c^{\text{tot}}}{e^{\beta\Delta F}\left(1+2e^{-\beta F/2}\right)^{1/2}\left(2+e^{\beta F/2}\right)^2} > \frac{\pi}{12}. \tag{S61}$$

###### S4.2.2 Monomers have (almost) all stickers typically bound

If $\frac{2n}{2+e^{\beta F/2}} \gtrsim n-2$, monomers typically have all or almost all their stickers bound. In that case, $Z_1$ is best approximated by that regime. However, for large enough $m$, $Z_m$ will be best approximated by the intermediate regime. This is because an approximately constant fraction of the stickers is typically bound, and for larger $m$, that fraction will correspond to a greater number of unbound stickers. We define $m^\star + 1$ to be the smallest value of $m$ for which the intermediate regime applies.

**Even $n$**

For even $n$, we found previously that $Z_1$ is approximately given by

$$Z_1 = e^{\left(\log(2)-\frac{\beta F}{2}\right)n}\left(\frac{1}{n+2}\sqrt{\frac{8}{n\pi}}\right). \tag{S62}$$

Since when $n$ is even, $nm$ is also always even, we have

$$Z_{m\leq m^\star} = e^{\left(\log(2)-\frac{\beta F}{2}\right)nm-\beta(m-1)\Delta F}\frac{1}{m}\left(\frac{1}{nm+2}\sqrt{\frac{8}{nm\pi}}\right). \tag{S63}$$

This gives a ratio

$$\frac{Z_{m\leq m^\star}}{Z_1^m} = m^{-3/2}\frac{n+2}{nm+2}\left[e^{-\beta\Delta F}(n+2)\sqrt{\frac{n\pi}{8}}\right]^{m-1} \tag{S64}$$

For $m > m^\star$, we have

$$Z_{m>m^\star} = \frac{e^{\beta\Delta F}\left(1+2e^{-\beta F/2}\right)^{1/2}}{2\pi m\left(\frac{nm}{2+e^{\beta F/2}}\right)\left(\frac{nm}{2+e^{\beta F/2}}+1\right)}\left[\left(1+2e^{-\beta F/2}\right)^n e^{-\beta\Delta F}\right]^m \tag{S65}$$

yielding the ratio

$$\frac{Z_{m>m^\star}}{Z_1^m} = \frac{e^{\beta\Delta F}\left(1+2e^{-\beta F/2}\right)^{1/2}}{2\pi m\left(\frac{nm}{2+e^{\beta F/2}}\right)\left(\frac{nm}{2+e^{\beta F/2}}+1\right)}\left[e^{-\beta\Delta F}\left(1+\frac{e^{\beta F/2}}{2}\right)^n(n+2)\sqrt{\frac{n\pi}{8}}\right]^m. \tag{S66}$$

We follow the same protocol as in the previous section, but our sum is now broken up into two parts:

$$\begin{aligned} c^{\text{tot}} &= \sum_{m=1}^{m^\star} m\frac{Z_{m\leq m^\star}}{Z_1^m}c_1^m + \sum_{m=m^\star+1}^{\infty} m\frac{Z_{m>m^\star}}{Z_1^m}c_1^m \\ &= e^{\beta\Delta F}\sqrt{\frac{8}{n\pi}}\sum_{m=1}^{m^\star}\frac{1}{(nm+2)\sqrt{m}}\left[y(c_1)\right]^m + \\ &\quad \frac{e^{\beta\Delta F}\left(1+2e^{-\beta F/2}\right)^{1/2}\left(2+e^{\beta F/2}\right)}{2\pi n}\sum_{m=m^\star+1}^{\infty}\frac{1}{m\left(\frac{nm}{2+e^{\beta F/2}}+1\right)}\left[y(c_1)\left(1+\frac{e^{\beta F/2}}{2}\right)^n\right]^m \end{aligned} \tag{S67}$$

where we have defined

$$y(c_1) = e^{-\beta\Delta F}(n+2)\sqrt{\frac{n\pi}{8}}\,c_1 \tag{S68}$$

If we make the approximation that $n \gg 2$ (for the first sum) and $\frac{nm^\star}{2+e^{\beta F/2}} \gg 1$ (for the second), the equations become a bit cleaner. Note that we could also not make such an approximation, and instead further split each sum in two; the first new sum (for smaller values of $m$) would then remain unapproximated and need to be computed explicitly, and the second (for larger values) will be approximated with the aforementioned approximations. For clarity and simplicity, we do not split the sums up further here. We thus arrive at the slighly simpler equation

$$c^{\text{tot}} = e^{\beta\Delta F}\sqrt{\frac{8}{n^3\pi}}\sum_{m=1}^{m^\star} m^{-3/2}\left[y(c_1)\right]^m + \frac{e^{\beta\Delta F}\left(1+2e^{-\beta F/2}\right)^{1/2}\left(2+e^{\beta F/2}\right)^2}{2\pi n^2}\sum_{m=m^\star+1}^{\infty} m^{-2}\left[y(c_1)\left(1+\frac{e^{\beta F/2}}{2}\right)^n\right]^m . \tag{S69}$$

The first sum will always be finite; aggregation is predicted when $y(c_1) > \left(1+\frac{e^{\beta F/2}}{2}\right)^{-n}$ (and therefore the second sum diverges). This inequality is substituted by an equality at the aggregation threshold itself, thus defining the value of $c_1$ at the threshold. By plugging in this value into the first sum, we find that the aggregation is predicted when $c^{\text{tot}} > c^{\text{tot}}_{\text{thresh}}$, defined by

$$c^{\text{tot}}_{\text{thresh}} = e^{\beta\Delta F}\sqrt{\frac{8}{n^3\pi}}\sum_{m=1}^{m^\star} m^{-3/2}\left[\left(1+\frac{e^{\beta F/2}}{2}\right)^{-n}\right]^m + \frac{e^{\beta\Delta F}\left(1+2e^{-\beta F/2}\right)^{1/2}\left(2+e^{\beta F/2}\right)^2}{2\pi n^2}\sum_{m=m^\star+1}^{\infty} m^{-2}. \tag{S70}$$

The result of the second sum is given by $\psi^{(1)}(m^\star+1)$ where $\psi$ is the polygamma function defined previously. The result of the first sum can either be written in terms of the Lerch transcendent $\Phi$ and polylogarithm function $Li_{3/2}$, or simply evaluated directly. The latter approach needs no further explanation, and for many purposes is the most straightforward approach to take. To demonstrate the former approach, we define the two functions here:

$$\begin{aligned}\Phi(z,s,\alpha) =& \sum_{k=0}^{\infty}\frac{z^k}{(k+\alpha)^s}\\ Li_s(z) =& \Phi(z,s,0) = \sum_{k=0}^{\infty}\frac{z^k}{k^s}\end{aligned} \tag{S71}$$

and use them to write

$$\sum_{m=1}^{m^\star} m^{-3/2}\left[\left(1+\frac{e^{\beta F/2}}{2}\right)^{-n}\right]^m = Li_{3/2}\left(\left(1+\frac{e^{\beta F/2}}{2}\right)^{-n}\right) - \left(1+\frac{e^{\beta F/2}}{2}\right)^{-n(1+m^\star)}\Phi\left(\left(1+\frac{e^{\beta F/2}}{2}\right)^{-n},\frac{3}{2},m^\star+1\right). \tag{S72}$$

The benefit of writing the sum in this way is that these two functions have been deeply categorized and explored mathematically (and that therefore, functions in Python, Mathematica, and similar programs can be used to evaluate them very efficiently). Putting it together, we find that we predict aggregation in this regime when

$$\frac{c^{\text{tot}}}{e^{\beta\Delta F}} > \frac{\left(1+2e^{-\beta F/2}\right)^{1/2}\left(2+e^{\beta F/2}\right)^2}{2\pi n^2}\psi^{(1)}(m^\star+1) + \sqrt{\frac{8}{n^3\pi}}\left[Li_{3/2}\left(\left(1+\frac{e^{\beta F/2}}{2}\right)^{-n}\right) - \left(1+\frac{e^{\beta F/2}}{2}\right)^{-n(1+m^\star)}\Phi\left(\left(1+\frac{e^{\beta F/2}}{2}\right)^{-n},\frac{3}{2},m^\star+1\right)\right]. \quad \text{(S73)}$$

A reasonable estimate of $m^\star$ is the value of $m$ for which $N_s(m^\star) = nm^\star - 2$, yielding

$$m^\star = \frac{2+4e^{-\beta F/2}}{n}. \quad \text{(S74)}$$

Plugging this result back into the previous equation we find a prediction for the concentration threshold that depends only on the values of $n$, $\beta F$, and $\beta\Delta F$.

**Odd $n$**

We follow the same procedure for odd values of $n$. However, now, $m < m^\star$, the value of $Z_m$ depends on whether $m$ is even or odd. We therefore now have three cases. Besides $Z_{m\le m^\star,\,\text{even}}$ and $Z_{m>m^\star}$ which are equivalent to the corresponding expressions for the even $n$ case, we have

$$\begin{aligned} Z_1 &= e^{\left(\log(2)-\frac{\beta F}{2}\right)(n-1)}\left(\frac{n}{n+1}\sqrt{\frac{8}{(n-1)\pi}}\right) \\ Z_{m\le m^\star,\,\text{odd}} &= e^{\left(\log(2)-\frac{\beta F}{2}\right)(nm-1)-\beta(m-1)\Delta F}\frac{1}{m}\left(\frac{nm}{nm+1}\sqrt{\frac{8}{(nm-1)\pi}}\right). \end{aligned} \quad \text{(S75)}$$

The three cases are

$$\begin{aligned} \frac{Z_{m\le m^\star,\,\text{odd}}}{Z_1^m} &= \left[2e^{-\frac{\beta F}{2}-\beta\Delta F}\left(\frac{n+1}{n}\right)\left(\frac{(n-1)\pi}{8}\right)^{1/2}\right]^m \frac{n}{nm+1}\left(\frac{8}{(nm-1)\pi}\right)^{1/2}\frac{e^{\frac{\beta F}{2}+\beta\Delta F}}{2} \\ \frac{Z_{m\le m^\star,\,\text{even}}}{Z_1^m} &= \left[2e^{-\frac{\beta F}{2}-\beta\Delta F}\left(\frac{n+1}{n}\right)\left(\frac{(n-1)\pi}{8}\right)^{1/2}\right]^m \frac{1}{m(nm+2)}\left(\frac{8}{nm\pi}\right)^{1/2}e^{\beta\Delta F} \\ \frac{Z_{m>m^\star}}{Z_1^m} &= \left[\left(1+\frac{e^{\beta F/2}}{2}\right)^n 2e^{-\frac{\beta F}{2}-\beta\Delta F}\left(\frac{n+1}{n}\right)\left(\frac{(n-1)\pi}{8}\right)^{1/2}\right]^m \frac{e^{\beta\Delta F}\left(1+2e^{-\beta F/2}\right)^{1/2}}{2\pi m\left(\frac{nm}{2+e^{\beta F/2}}\right)\left(\frac{nm}{2+e^{\beta F/2}}+1\right)} \end{aligned} \quad \text{(S76)}$$

In keeping with the previous procedure, we define $z(c_1)$ to be

$$z(c_1) = 2e^{-\frac{\beta F}{2}-\beta\Delta F}\left(\frac{n+1}{n}\right)\left(\frac{(n-1)\pi}{8}\right)^{1/2}c_1. \quad \text{(S77)}$$

The total concentration is then given by

$$
\begin{aligned}
c^{\text{tot}} = & \sum_{m=1,3,5\ldots}^{m^\star} \frac{nm}{nm+1}\left(\frac{8}{(nm-1)\pi}\right)^{1/2} \frac{e^{\frac{\beta F}{2}+\beta\Delta F}}{2}\left[z(c_1)\right]^m + \\
& \sum_{m=2,4,6\ldots}^{m^\star} \frac{1}{nm+2}\left(\frac{8}{nm\pi}\right)^{1/2} e^{\beta\Delta F}\left[z(c_1)\right]^m + \\
& \sum_{m=m^\star+1}^{\infty} \frac{e^{\beta\Delta F}\left(1+2e^{-\beta F/2}\right)^{1/2}}{2\pi\left(\frac{nm}{2+e^{\beta F/2}}\right)\left(\frac{nm}{2+e^{\beta F/2}}+1\right)}\left[\left(1+\frac{e^{\beta F/2}}{2}\right)^n z(c_1)\right]^m . \quad \text{(S78)}
\end{aligned}
$$

As previously, aggregation is predicted when the final sum diverges, or when $z(c_1) > \left(1+\frac{e^{\beta F/2}}{2}\right)^{-n}$. In order to simplify the mathematics, we make the approximation that $n \gg 2$. This allows us to write each summand as $m^{-\lambda}\gamma^m$ for some values of $\lambda$ and $\gamma$:

$$
\begin{aligned}
c^{\text{tot}} = & \frac{e^{\frac{\beta F}{2}+\beta\Delta F}}{2}\sqrt{\frac{8}{n\pi}} \sum_{m=1,3,5\ldots}^{m^\star} m^{-1/2}\left[z(c_1)\right]^m + \sqrt{\frac{8}{n^3\pi}} e^{\beta\Delta F} \sum_{m=2,4,6\ldots}^{m^\star} m^{-3/2}\left[z(c_1)\right]^m + \\
& \frac{e^{\beta\Delta F}\left(1+2e^{-\beta F/2}\right)^{1/2}}{2\pi\left(\frac{n}{2+e^{\beta F/2}}\right)^2} \sum_{m=m^\star+1}^{\infty} m^{-2}\left[\left(1+\frac{e^{\beta F/2}}{2}\right)^n z(c_1)\right]^m . \quad \text{(S79)}
\end{aligned}
$$

To address the first sum, we let $m_o = (m+1)/2$, such that

$$
\begin{aligned}
\sum_{m=1,3,5\ldots}^{m^\star} m^{-1/2}\left[z\right]^m &= \frac{1}{z\sqrt{2}} \sum_{m_o=1}^{\frac{m^\star+1}{2}} \left(m_o - \frac{1}{2}\right)^{-1/2}\left[z^2\right]^{m_o} \\
&= \frac{z}{\sqrt{2}}\left[\Phi\left(z^2, \frac{1}{2}, \frac{1}{2}\right) - z^{m^\star+1}\Phi\left(z^2, \frac{1}{2}, \frac{m^\star}{2}+1\right)\right] \quad \text{(S80)}
\end{aligned}
$$

(where we have written $z(c_1)$ as $z$ for notational convenience and clarity). For notational clarity, we will denote this combination by $s_o(z, m^\star)$. We make a similar substitution for the second sum (which we will denote by $s_e(z, m^\star)$), letting $m_e = m/2$:

$$
\begin{aligned}
\sum_{m=2,4,6\ldots}^{m^\star} m^{-3/2}\left[z\right]^m &= 2^{-3/2} \sum_{m_e=1}^{\frac{m^\star}{2}} m_e^{-3/2}\left(z^2\right)^{m_e} \\
&= 2^{-3/2}\left[Li_{3/2}(z^2) - z^{m^\star+2}\Phi\left(z^2, \frac{3}{2}, \frac{m^\star}{2}+1\right)\right]. \quad \text{(S81)}
\end{aligned}
$$

The aggregation threshold occurs when $z(c_1) = \left(1+\frac{e^{\beta F/2}}{2}\right)^{-n}$, meaning that

$$
\begin{aligned}
\frac{c^{\text{tot}}_{\text{thresh}}}{e^{\beta\Delta F}} = & e^{\frac{\beta F}{2}}\sqrt{\frac{2}{n\pi}}\, s_o\left(\left(1+\frac{e^{\beta F/2}}{2}\right)^{-n}, m^\star\right) + \\
& \sqrt{\frac{8}{n^3\pi}}\, s_e\left(\left(1+\frac{e^{\beta F/2}}{2}\right)^{-n}, m^\star\right) + \\
& \frac{\left(1+2e^{-\beta F/2}\right)^{1/2}}{2\pi\left(\frac{n}{2+e^{\beta F/2}}\right)^2}\psi^{(1)}(m^\star+1). \quad \text{(S82)}
\end{aligned}
$$

Given our previous estimate of $m^\star = \frac{2+4e^{-\beta F/2}}{n}$, this expression only depends on the parameters $n$, $\beta F$, and $\beta\Delta F$, and is therefore our final expression for the aggregation threshold in this regime.

#### S4.3 Disallowing neighbor bonds

The procedure here follows that outlined in the previous section, with the relevant partition functions substituted for one another. Because the procedure is so similar, we will move faster through these calculations.

As we saw, the partition function for multimers in the intermediate regime appears in all of the calculations. If we define a function $f(\beta F)$ such that

$$f(\beta F) = 1 - \frac{e^{\beta F/4}}{\sqrt{4 + e^{\beta F/2}}} \tag{S83}$$

then the dominant term in that partition function is that corresponding to $N_b^\star = qf(\beta F)/2$. Recognizing that in the intermediate binding regime, $N_b^\star \gg 1$, the equation for $Z_m$ in this regime (considering only the dominant term) is

$$Z_m = \frac{2}{\pi} \left[qf(\beta F)\right]^{-2} \frac{e^{-\beta(m-1)\Delta F}}{m} \left( \left[ 2e^{-\beta F/2} \left( \frac{1 - f(\beta F)}{1 - \frac{f(\beta F)}{2}} \right) \left( \frac{1}{f(\beta F)} - 1 \right) \right]^{f(\beta F)} \left[ \frac{1 - \frac{f(\beta F)}{2}}{1 - f(\beta F)} \right]^2 \right)^q . \tag{S84}$$

For clarity, we denote by $\mathcal{F}(\beta F)$ the expression raised to the power of $q$, such that the expression above can be written a bit more cleanly:

$$Z_m = \frac{2}{\pi} \left[qf(\beta F)\right]^{-2} \frac{e^{-\beta(m-1)\Delta F}}{m} \left[\mathcal{F}(\beta F)\right]^q . \tag{S85}$$

##### S4.3.1 Monomers have some stickers typically bound

In this regime, we have

$$\frac{Z_m}{Z_1^m} = \frac{n^2}{mq^2} \left[ \frac{\pi}{2} \left(f(\beta F)\right)^2 n^2 e^{-\beta \Delta F} \left(\mathcal{F}(\beta F)\right)^\alpha \right]^{m-1} . \tag{S86}$$

where $q = nm + \alpha(m-1)$. If we approximate the $q$ in the denominator as $m(n+\alpha)$ (assuming that $nm \gg \alpha$, a reasonable assumption given that $\alpha < 1$), the prefactor becomes

$$\frac{1}{m^3} \frac{n^2}{(n+\alpha)^2} . \tag{S87}$$

Defining

$$x(c_1) = \frac{\pi}{2} \left(f(\beta F)\right)^2 n^2 e^{-\beta \Delta F} \left(\mathcal{F}(\beta F)\right)^\alpha c_1 \tag{S88}$$

we have

$$c^{\text{tot}} \frac{(n+\alpha)^2}{n^2} x(1) = \sum_{m=1}^{\infty} \frac{1}{m^2} \left[x(c_1)\right]^m . \tag{S89}$$

We therefore predict that aggregation occurs when

$$\left(f(\beta F)\right)^2 \left(\mathcal{F}(\beta F)\right)^\alpha (n+\alpha)^2 \frac{c^{\text{tot}}}{e^{\beta \Delta F}} > \frac{\pi}{3} . \tag{S90}$$

##### S4.3.2 Monomers have (almost) all stickers typically bound

In the corresponding section in which we allowed neighbor bonds, we split up the respective sums into those for $m \le m^\star$ and $m > m^\star$, with $m^\star$ defined as the value of $m$ for which the multimer partition function begins to be better approximated by the intermediate regime than by the regime in which all stickers are typically bound. We could certainly do the same split here; however, we find that typical values of $m^\star$ are so small (typically 3), and the intermediate regime such a good approximation, as to make it reasonable to forgo this split. We instead simply consider all multimers with $m > 2$ as being well-approximated by the intermediate regime. Nevertheless, if a different split is needed, it follows along the same lines as when allowing neighbor bonds (with the exception that even and odd $m$ should to be considered separately for all $n$ and not only for odd $n$).

**Even** $n$
For even $n$, the dominant term in the monomer partition function is

$$Z_1 = \frac{n(n+2)}{8} e^{-\frac{\beta F}{2}(n-2)} \tag{S91}$$

Thus, for large $m$,

$$\frac{Z_{m>m^\star}}{Z_1^m} = \frac{2}{\pi}\left(\frac{1}{nm+\alpha(m-1)}\right)^2 (f(\beta F))^{-2}\frac{e^{\beta\Delta F}}{m}(\mathcal{F}(\beta F))^{-\alpha}\left((\mathcal{F}(\beta F))^{n+\alpha} e^{-\beta\Delta F}\frac{8}{n(n+2)}e^{\frac{\beta F}{2}(n-2)}\right)^m. \tag{S92}$$

For small $m$,

$$\frac{Z_{m\leq m^\star}}{Z_1^m} = e^{\beta F\frac{\alpha}{2}}\frac{e^{\beta\Delta F}}{m}\left[e^{-\beta F(1+\alpha/2)}\left(\frac{8}{n(n+2)}\right)e^{-\beta\Delta F}\right]^m. \tag{S93}$$

Approximating $m^\star(n+\alpha) \gg \alpha$ (i.e. $n \gg \alpha/m^\star$), the total concentration is

$$\begin{aligned} c^{\text{tot}} &= \sum_{m=1}^{m^\star} m\frac{Z_{m\leq m^\star}}{Z_1^m}c_1^m + \sum_{m=m^\star+1}^{\infty} m\frac{Z_{m>m^\star}}{Z_1^m}c_1^m \\ &= e^{\beta F\frac{\alpha}{2}}e^{\beta\Delta F}\sum_{m=1}^{m^\star}\left[y(c_1)\left(\mathcal{F}(\beta F)e^{\frac{\beta F}{2}}\right)^{-n-\alpha}\right]^m + \frac{2(\mathcal{F}(\beta F))^{-\alpha}e^{\beta\Delta F}}{\pi(n+\alpha)^2(f(\beta F))^2}\sum_{m=m^\star+1}^{\infty}\frac{1}{m^2}[y(c_1)]^m. \end{aligned} \tag{S94}$$

where we have defined

$$y(c_1) = (\mathcal{F}(\beta F))^{n+\alpha} e^{-\beta\Delta F}\frac{8}{n(n+2)}e^{\frac{\beta F}{2}(n-2)}c_1. \tag{S95}$$

The aggregation transition is defined by $y(c_1) = 1$, or $c_1 = 1/y(1)$. Thus, aggregation is predicted when

$$c^{\text{tot}} > e^{\beta F\frac{\alpha}{2}}e^{\beta\Delta F}\sum_{m=1}^{m^\star}\left[\left(\mathcal{F}(\beta F)e^{\frac{\beta F}{2}}\right)^{-(n+\alpha)}\right]^m + \frac{2(\mathcal{F}(\beta F))^{-\alpha}e^{\beta\Delta F}}{\pi(n+\alpha)^2(f(\beta F))^2}\psi^{(1)}(m^\star+1). \tag{S96}$$

The first sum can also be calculated directly, yielding

$$c^{\text{tot}} > e^{\beta F\frac{\alpha}{2}}e^{\beta\Delta F}\frac{\left(\mathcal{F}(\beta F)e^{\frac{\beta F}{2}}\right)^{-(n+\alpha)}\left[1-\left(\mathcal{F}(\beta F)e^{\frac{\beta F}{2}}\right)^{-(n+\alpha)m^\star}\right]}{1-\left(\mathcal{F}(\beta F)e^{\frac{\beta F}{2}}\right)^{-(n+\alpha)}} + \frac{2(\mathcal{F}(\beta F))^{-\alpha}e^{\beta\Delta F}}{\pi(n+\alpha)^2(f(\beta F))^2}\psi^{(1)}(m^\star+1). \tag{S97}$$

A reasonable estimate of $m^\star$ is the value of $m$ for which $N_s(m^\star) = nm^\star - 2$, yielding

$$m^\star = \frac{2}{n(1-f(\beta F))} = \frac{2\sqrt{4+e^{\beta F/2}}}{ne^{\beta F/4}}. \tag{S98}$$

Plugging this result back into the previous equation we find a prediction for the concentration threshold that depends only on the values of $n$, $\beta F$, and $\beta\Delta F$.

As discussed, for the free energies we consider, we typically find $m^\star = 2$. We will now redo the calculation for that particular value. If we set $m^\star = 2$, we can separate out the $m = 1$ and $m = 2$ terms from the sum. We then have

$$
\begin{aligned}
c^{\text{tot}} - c_1 - 2\frac{Z_2}{Z_1^2}c_1^2 &= \sum_{m=3}^{\infty} m\frac{Z_m}{Z_1^m}c_1^m \\
&= \frac{2}{\pi}\left(f(\beta F)\right)^{-2} e^{\beta\Delta F}\left(\mathcal{F}(\beta F)\right)^{-\alpha}\sum_{m=3}^{\infty}\left(\frac{1}{(n+\alpha)m-\alpha}\right)^2\left[y(c_1)\right]^m
\end{aligned}
\tag{S99}
$$

where

$$
2\frac{Z_2}{Z_1^2} = \left(\frac{8}{n(n+2)}\right)^2 e^{-\beta\Delta F - \beta F(2+\alpha/2)}\left(1+\mathcal{O}(e^{\beta F})\right) \tag{S100}
$$

and we have defined the same $y(c_1)$ as for the general $m^\star$ case.

$$
y(c_1) = \left(\mathcal{F}(\beta F)\right)^{n+\alpha} e^{-\beta\Delta F}\frac{8}{n(n+2)}e^{\frac{\beta F}{2}(n-2)}c_1. \tag{S101}
$$

Approximating $3(n+\alpha) \gg \alpha$ (i.e. $n \gg \alpha/3$), the $(n+\alpha)^2$ term can be taken out of the sum.

Since the aggregation transition is defined by $y(c_1) = 1$, or $c_1 = 1/y(1)$, aggregation is predicted when

$$
\begin{aligned}
&\left[(n+\alpha)f(\beta F)\right]^2\Bigg[\left(\mathcal{F}(\beta F)\right)^{\alpha}\frac{c^{\text{tot}}}{e^{\beta\Delta F}} - \\
&\qquad \frac{n(n+2)}{8}\left(\mathcal{F}(\beta F)\right)^{-n}e^{-\frac{\beta F}{2}(n-2)} - \left(\mathcal{F}(\beta F)\right)^{-2n-\alpha}e^{-\beta F(n+\alpha/2)}\Bigg] > \frac{\pi}{3} - \frac{5}{2\pi}.
\end{aligned}
\tag{S102}
$$

**Odd** $n$

For odd $n$, the dominant term in the monomer partition function is

$$
Z_1 = e^{-\frac{\beta F}{2}(n-1)}. \tag{S103}
$$

The approach follows the same procedure as for even $n$. Instead of $y(c_1)$ though, we have

$$
z(c_1) = \left(\mathcal{F}(\beta F)\right)^{n+\alpha} e^{-\beta\Delta F} e^{\frac{\beta F}{2}(n-1)}c_1. \tag{S104}
$$

We also have three cases to consider:

$$
\begin{aligned}
\frac{Z_{m>m^\star}}{Z_1^m} &= \frac{2}{\pi}\left(f(\beta F)\right)^{-2} e^{\beta\Delta F}\left(\mathcal{F}(\beta F)\right)^{-\alpha}(n+\alpha)^{-2}\frac{1}{m^3}z(1)^m \\
\frac{Z_{m\le m^\star,\,\text{odd}}}{Z_1^m} &= \frac{e^{\beta\Delta F}}{m}e^{\frac{\beta F}{2}(1+\alpha)}\left[e^{-\beta\Delta F}e^{-\frac{\beta F}{2}(1+\alpha)}\right]^m \\
\frac{Z_{m\le m^\star,\,\text{even}}}{Z_1^m} &= \frac{e^{\beta\Delta F}}{m}e^{\frac{\beta F}{2}\alpha}\left[e^{-\beta\Delta F}e^{-\frac{\beta F}{2}(1+\alpha)}\right]^m.
\end{aligned}
\tag{S105}
$$

The total concentration is

$$
\begin{aligned}
c^{\text{tot}} = e^{\beta\Delta F}e^{\frac{\beta F}{2}(1+\alpha)} &\sum_{m=1,3,5\ldots}^{m^\star}\left[\left(\mathcal{F}(\beta F)\right)^{-(n+\alpha)}e^{-\frac{\beta F}{2}(n+\alpha)}z(c_1)\right]^m + \\
&e^{\beta\Delta F}e^{\frac{\beta F}{2}\alpha}\sum_{m=2,4,6\ldots}^{m^\star}\left[\left(\mathcal{F}(\beta F)\right)^{-(n+\alpha)}e^{-\frac{\beta F}{2}(n+\alpha)}z(c_1)\right]^m + \\
&\qquad \frac{2}{\pi}\left(f(\beta F)\right)^{-2}e^{\beta\Delta F}\left(\mathcal{F}(\beta F)\right)^{-\alpha}(n+\alpha)^{-2}\sum_{m=m^\star+1}^{\infty}\frac{1}{m^2}z(c_1)^m.
\end{aligned}
\tag{S106}
$$

Aggregation is predicted when $z(c_1) > 1$, or

$$c^{\text{tot}} > e^{\beta\Delta F} e^{\frac{\beta F}{2}\alpha} \frac{w\left(1 + e^{\beta F/2} - w^{\frac{m^\star}{2}}\left[1 + w^{1/2} e^{\beta F/2}\right]\right)}{1-w} + \frac{2}{\pi}\left(f(\beta F)\right)^{-2} e^{\beta\Delta F}\left(\mathcal{F}(\beta F)\right)^{-\alpha}(n+\alpha)^{-2}\psi^{(1)}(m^\star + 1) \quad \text{(S107)}$$

where

$$w = \left(\mathcal{F}(\beta F) e^{\frac{\beta F}{2}}\right)^{-(n+\alpha)}. \quad \text{(S108)}$$

Setting $m^\star$ to (S98), this result depends only on the values of $n$, $\beta F$, and $\beta\Delta F$.

With this result in hand, we can also do the calculation setting $m^\star = 2$. For odd $n$, the dimer term yields

$$2\frac{Z_2}{Z_1^2} = e^{-\beta\Delta F - \beta F(1+\alpha/2)}\left(1 + \mathcal{O}(e^{\beta F})\right). \quad \text{(S109)}$$

With the substitution of $z$ for $y$, the equation for $c^{\text{tot}}$ is the same as for even $n$:

$$c^{\text{tot}} - c_1 - 2\frac{Z_2}{Z_1^2}c_1^2 = \frac{2}{\pi}\left(f(\beta F)\right)^{-2} e^{\beta\Delta F}\left(\mathcal{F}(\beta F)\right)^{-\alpha}(n+\alpha)^{-2}\sum_{m=3}^{\infty}\frac{1}{m^2}\left[z(c_1)\right]^m \quad \text{(S110)}$$

The aggregation threshold is in this case defined by $c_1 = 1/z(1)$, such that aggregation is predicted when

$$\left[(n+\alpha) f(\beta F)\right]^2\left[\left(\mathcal{F}(\beta F)\right)^{\alpha}\frac{c^{\text{tot}}}{e^{\beta\Delta F}} - \left(\mathcal{F}(\beta F)\right)^{-n} e^{-\frac{\beta F}{2}(n-1)} - \left(\mathcal{F}(\beta F)\right)^{-2n-\alpha} e^{-\beta F(n+\alpha/2)}\right] > \frac{\pi}{3} - \frac{5}{2\pi}. \quad \text{(S111)}$$

##### S4.3.3 Monomers have almost no stickers bound

Because of the computational difficulty of computing partition functions in this regime, and the lack of a simple analytical formula for $g$, we do not consider it here. We believe this regime – in which monomers have almost no stickers bound, but the multiplicity of possible binding combinations drives binding for multimers – to be an interesting potential area for future research.

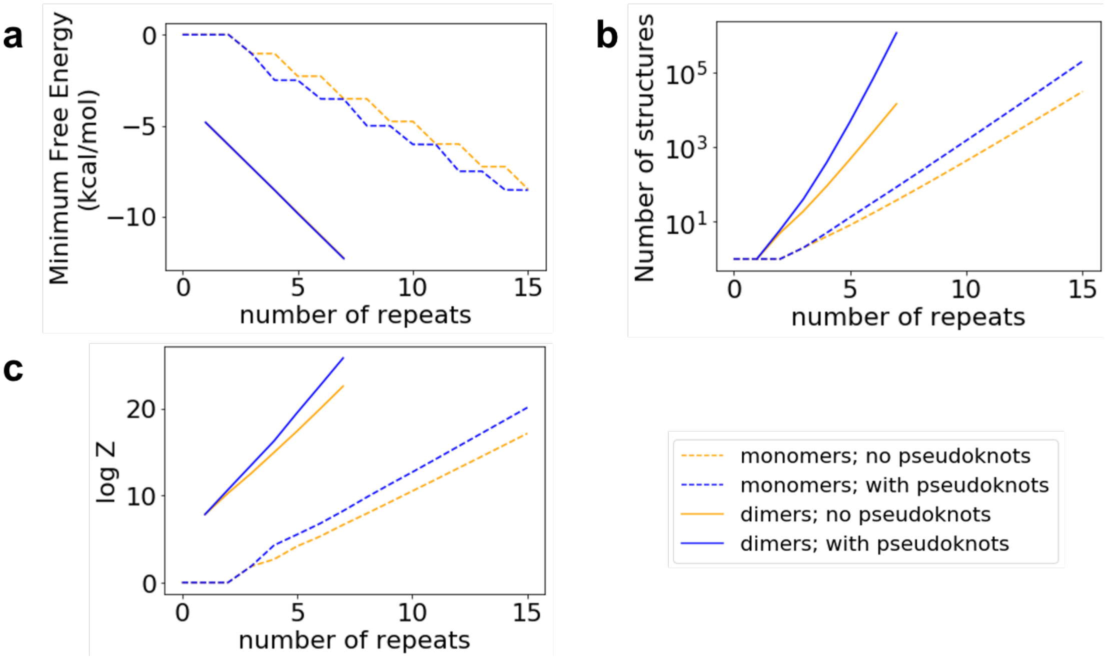

Figure S4: **Repeat RNA pseudoknots**. We consider the landscape of structures formed by RNA molecules consisting of $n$ CAG repeats for values of $n \leq 15$ (monomers; dashed) or $n \leq 7$ (dimers; solid lines). We either disallow (orange) or allow (blue) both intra- and inter-molecular pseudoknots. **a:** The minimum free energy (MFE) structure for monomers is pseudoknotted. The energy gap between the pseudoknotted and the non-pseudoknotted MFE structure is between $\sim 1.25 - 1.5$ kcal/mol for even values of $n$ (and smaller for odd $n$) but does not appear to grow with $n$. The dimeric MFE structure is always non-pseudoknotted. **b:** The number of structures enumerated grows significantly when allowing pseudoknots. For $n = 7$ dimers, $\sim 14,000$ non-pseudoknotted structures are enumerated compared to $> 1.1$ million pseudoknotted structures. **c:** The partition function is affected by the inclusion or exclusion of pseudoknotted structures, but this effect does not appear particularly significant for our purposes (see Fig. S5).

### S5 Computational enumeration

#### S5.1 Complete enumeration is used to probe the contribution of pseudoknots to $Z$

How much do pseudoknots affect the landscape of structures? To answer this question, we used LandscapeFold [9] to enumerate all monomeric and dimeric structures that can form with $n$ CAG repeats ($n \leq 15$ for monomeric structures; $n \leq 7$ for dimeric structures). The results are shown in Fig. S4.

In panel A we show that the minimum free energy (MFE) structure is a pseudoknot for monomeric structures for $n \geq 4$. However, for odd $n$, its free energy is almost equal to the non-pseudoknotted MFE structure. Moreover, the energy gap between MFE pseudoknotted and non-pseudoknotted structures appears constant as a function of $n$ (aside from the even/odd discrepancy). For dimers, the MFE structure is always non-pseudoknotted.

More significant is the effect of including pseudoknots on the landscape as a whole. In panel B we show that the number of pseudoknotted structures vastly outweighs the number of non-pseudoknotted for large $n$, especially for dimers. This multiplicity affects the partition function calculation (panel C). While all four partition functions here grow roughly exponentially with the number of repeats, the slope of the exponentials is higher when accounting for pseudoknots.

To quantify the effect of disallowing pseudoknots on our model results, we consider the predicted yields of the dimers while allowing and disallowing pseudoknots. We refer to the ratio $Z_2/Z_1^2$ as $r$ here. We define the relative error in $r$ due to disallowing pseudoknots as $(\log(r_n) - \log(r_p))/\log(r_p)$ where $r_n$ describes the results of the calculation disallowing pseudoknots, and $r_p$ the results allowing pseudoknots. Fig. S5 shows how the relative error changes as a function of the number of repeats $n$ in the RNA. We find that the error is mostly within $\sim 10\%$ (aside from $n = 4$ which has a $\sim 25\%$ error). Especially, we find that the error does not appear to grow with longer repeat lengths: the

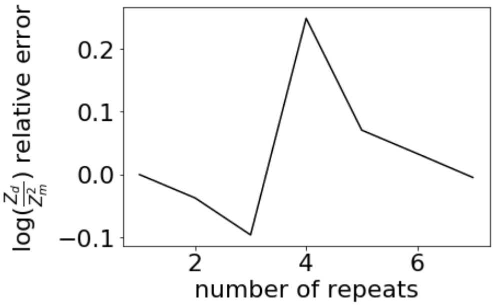

Figure S5: **The relative error due to disallowing pseudoknots**. We consider the landscapes of structures formed by RNA molecules consisting of $n$ CAG repeats. As in Fig. S4, we consider two landscapes: one in which we disallow pseudoknots; the other which includes them in the enumeration. We quantify the relative error of the ratio $Z_2/Z_1^2$ due to disallowing pseudoknots as described in the main text, and show the results as a function of $n$. We find that the relative error lies mostly within $10\%$ and does not appear to grow with $n$. In fact, the relative error due to disallowing pseudoknots for the longest RNA we considered here, $n = 7$, is $< 1\%$. We therefore do not consider pseudoknotted structures further in our model.

error for the longest RNA we considered with pseudoknots, $n = 7$, is $< 1\%$. The effects of disallowing pseudoknots therefore do not appear particularly significant for our purposes, and we do not consider pseudoknots further.

#### S5.2 Description of the dynamic programming algorithm

##### S5.2.1 General notes on the procedure

In this section we describe how we calculate $Z_m$ computationally by enumerating all possible structures that an arbitrary set of strands can form. By neglecting pseudoknots, we can employ a dynamic programming methodology that allows us to perform the enumeration in polynomial time (although the enumeration of certain classes of pseudoknots using dynamic programming approaches is possible; see e.g. Ref. [10]).

We exactly calculate the loop entropies for each structure. Non-pseudoknotted loops of length $s$ have an entropic penalty given by [11, 9]

$$\Delta S_{\text{loop}}(s) = k_B \left[ \ln v_s + \frac{3}{2} \ln \left( \frac{\gamma}{\pi s} \right) \right] \tag{S112}$$

where $v_s = 0.02$ nucleotides$^3$ is the volume within which two nucleotides can bind, and $\gamma = 3/2b$ where $b$ is the persistence length of single-stranded RNA (we used $b = 2.4$ nucleotides here for consistency with Ref. [9]). The length of the loop $s$ is calculated by taking the number of phosphodiester bonds in the loop. For example, hairpin loops comprised of $s$ nucleotides have a length of $s + 1$, while internal loops comprised of $s$ nucleotides have a length of $s + 2$.

Let us first consider monomers. Given the no-pseudoknot approximation, if a binding event between sites $i$ and $j$ is present, all the sites between $i + 1$ and $j - 1$ can only pair to one another, and therefore can be treated like an RNA molecule comprised of $j - i - 1$ binding sites. If another bond is formed between nucleotides $i' > i$ and $j' < j$, there are two loops that form: one of length $j' - i'$, and one of length $j' - i'$ and the other of length $i' - i + j' - j$. Thus, the length of the loop formed by the bond $(i, j)$ depends on the structure in the middle $j - i - 1$ nucleotides. For example, if $i = 5$ and $j = 10$, if there are no binding events in those middle binding sites, the length of the loop will be 14 nucleotides long, while if there is a binding event between sites 6 and 8, the loop will only be 7 nucleotides long.

To keep track of this feature, we define for each structure a quantity we call the phantom "outer loop": how long would a loop be that connects a binding site at position 0 with a binding site at position $n + 1$? This phantom outer loop does not enter into the free energy calculation for $n$ repeats, but does enter into the calculation when we use the results from $n$ repeats to calculate the landscape of a longer RNA with this number of repeats in the "middle" section.

Moving on to multimers, we consider a set of $m$ RNA strands, each comprised of $n_i$ CAG repeats (with $i$ ranging from 1 to $m$). The procedure is effected by calculating the landscape for the set of sequences of lengths ($n_1$, $n_2$, ...,

$n_m - 1$), and then adding to that the ensembles of structures that can form for each possible binding event involving the final binding site.

##### S5.2.2 The recursive relation underlying the algorithm

We keep track of three quantities: (1) $Z(n_1, n_2, ..., n_m)$, the partition function for the set of sequences being considered; (2) $Z_s(n_1, n_2, ..., n_m)$, the partition function of the ensemble of structures with a phantom outer loop of length $s$; (3) $\tilde{Z}(n_1, n_2, ..., n_m)$, the value taken by $Z$ when the entropy costs of the various phantom outer loops are taken into account. These three quantities are defined as:

$$Z_s(n_1, n_2, ..., n_m) = \sum_{\sigma_s} \exp(-\beta F_{\sigma_s}) \tag{S113}$$

$$Z(n_1, n_2, ..., n_m) = \sum_{\sigma} \exp(-\beta F_{\sigma}) = \sum_{s} Z_s(n_1, n_2, ..., n_m) \tag{S114}$$

$$\tilde{Z}(n_1, n_2, ..., n_m) = \sum_{\sigma} e^{-\beta(F_\sigma + F_b - T\Delta S_{\text{loop}}(s_\sigma))} = \sum_{s} Z_s(n_1, n_2, ..., n_m) e^{-\beta F_b + \Delta S_{\text{loop}}(s)/k_B} \tag{S115}$$

where $\sigma$ is a structure defined by a set of binding events, $F_\sigma$ is its free energy and $s_\sigma$ is its phantom outer loop length; $\sigma_s$ is a structure with a phantom outer loop of length $s$ and $F_{\sigma_s}$ is its free energy. As can be seen from these equations, $Z$ and $\tilde{Z}$ can both be written in terms of $Z_s$.

Extending our previous definition of the phantom outer loop, for a set of strands it is defined as the loop formed by a binding event between a binding site at position 0 of the first sequence and position $n_m + 1$ of the final sequence. If no closed loop is formed by such a binding event (e.g. for a dimer if there are no other intermolecular binding events) there is no explicit entropy cost to the phantom outer loop forming ($\Delta S_{\text{loop}} = 0$). Instead, the penalty will be given by $\Delta G_{\text{assoc}}$; we will add in appropriate factors of $\Delta G_{\text{assoc}}$ later in this section. For such phantom outer loops, it is useful to consider their lengths to be infinite in the following formulae (despite the fact that $\Delta S_{\text{loop}}(\infty) \neq 0$). This allows us in formulae which have terms such as $s - 2$ and $s - 3$ to consider these phantom outer loops to be unchanged by such subtractions.

We first consider the ensemble of structures in which the last binding site is unbound, and then consider each possible site $i$ to which the last binding site can bind. When the final binding site is bound to a site $i$, the set of RNA molecules is effectively split in two parts, since by disallowing pseudoknots we disallow any binding events between sites to the right of $i$ and sites to the left. For multimers of $m \geq 3$, some non-pseudoknotted structures are disallowed by this assumption; however, as discussed in Section S1.2.3, this actually works to our benefit since these are always identical to structures previously enumerated and we ultimately wish to enumerate each structure only once.

We describe our sum over binding sites $i$ as the combination of a sum over the strand $m'$ and a separate sum over the binding sites $i'$ in that strand. We also consider the case of $m' = m$ separately since when disallowing neighbor bonds, the final site cannot bind to the one immediately preceding it (because of constraints on hairpin loop length).

$$\begin{aligned} Z_s(n_1, n_2, ..., n_m + 1) =& Z_{s-3}(n_1, n_2, ..., n_m) + \\ & \sum_{m'=1}^{m-1} \sum_{i'=1}^{n_i} Z_{s-2}(n_1, n_2, ..., n_{m'-1}, i' - 1) \tilde{Z}(n_{m'} - i', n_{m'+1}, ..., n_m) + \\ & \sum_{i'=1}^{n_m - 1} Z_{s-2}(n_1, n_2, ..., n_{m-1}, i' - 1) \tilde{Z}(n_m - i). \end{aligned} \tag{S116}$$

Here, the last sum includes $n_m$ if allowing neighbor bonds. We also require the base cases: $Z_\infty(n_1, n_2, ..., n_{m-1}, 0) = Z(n_1, n_2, ..., n_{m-1})$; $Z_2(0) = Z_5(1) = Z_8(2) = 1$ (the last case is not included when allowing neighbor bonds).

With this procedure, we are in principle able to calculate the landscapes of arbitrary multimers. The scaling of compute time with repeat length is shown in Fig. S6. Given the recursive nature of the algorithm, we display only the additional computation time necessary to compute Eqns. S113-S115 for a set of $m$ strands comprised of $n_i = n$ repeats given that the landscapes for a set of $m$ strands comprised of $n_i = n - 1$ repeats and for a set of $m - 1$ strands comprised of $n$ repeats have both already been computed.

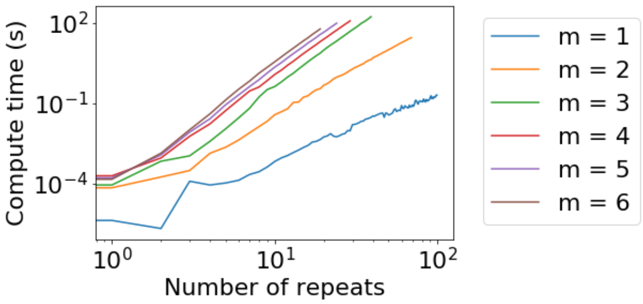

Figure S6: **Compute time for multimer landscape calculation.** The computation times for multimers of $m$ strands each comprised of $n$ repeats is shown. Given the recursive nature of the algorithm, we display only the additional computation time necessary to compute $Z$ for a multimer comprised of $n$ repeats given that the landscape for a multimer comprised of $n-1$ repeats has already been computed.

##### S5.2.3 Computing $Z$ for single complexes and correcting for symmetries

In the previous subsections, we showed how to compute the landscape of all structures that can be formed by a set of $m$ strands of different lengths. However, that landscape includes both $m$-mers as well as monomers and $m-1$-mers, dimers and $m-2$-mers, etc. In this subsection we consider how to use these landscapes to compute $Z$ for single complexes comprised of $m$ strands. This is analogous to the correction described in Section S1.2.4. We also correct for symmetries as discussed in Section S1.2.3.

For $m = 2$, there are two types of structures: dimers and pairs of monomers. Therefore,

$$Z_2(n_1, n_2) = \frac{1}{2}\Big(Z(n_1, n_2) - Z(n_1)Z(n_2)\Big). \tag{S117}$$

The factor of $1/2$ corrects for that every (asymmetric) dimer structure is counted twice in $Z(n_1, n_2)$ (see Section S1.2.3). The subtraction accounts for that $Z(n_1, n_2)$ includes not only dimers but pairs of monomers as well.

For $m = 3$, there are three types of structures: trimers, 3 monomers, and 1 monomer and 1 dimer. We therefore have

$$Z_3(n_1, n_2, n_3) = \frac{1}{3}Z(n_1, n_2, n_3) - \frac{1}{3}\Big(Z(n_1)Z(n_2)Z(n_3) + 2Z_2(n_1, n_2)Z(n_3) + 2Z_2(n_1, n_3)Z(n_2) + 2Z_2(n_2, n_3)Z(n_1)\Big). \tag{S118}$$

The factor of 2 before each factor of $Z_2$ accounts for the fact that $Z(n_1, n_2, n_3)$ overcounted the dimer & monomer structures in the same way $Z_2$ did (i.e. by a factor of 2). The factor of 2 here ensures we properly subtract the contribution of dimers & monomers from $Z(n_1, n_2, n_3)$.

For $m = 4$ the calculation is somewhat complicated by our no-pseudoknot assumption. The system can form a tetramer, 4 monomers, 1 trimer and 1 monomer, 2 monomers and 1 dimer, or 2 dimers. However not every set of two dimers can form, since the pair of dimers $(n_1, n_3), (n_2, n_4)$ looks like a pseudoknot in our model and was therefore not enumerated. Our model therefore yields

$$\begin{aligned} Z_4(n_1, n_2, n_3, n_4) = \frac{1}{4}Z(n_1, n_2, n_3, n_4) - \frac{1}{4}\Big(&Z(n_1)Z(n_2)Z(n_3)Z(n_4) + \\ &3Z_3(n_1, n_2, n_3)Z(n_4) + 3Z_3(n_1, n_2, n_4)Z(n_3) + 3Z_3(n_1, n_3, n_4)Z(n_2) + \\ &3Z_3(n_2, n_3, n_4)Z(n_1) + 2Z_2(n_1, n_2)Z(n_3)Z(n_4) + 2Z_2(n_1, n_3)Z(n_2)Z(n_4) + \\ &2Z_2(n_1, n_4)Z(n_2)Z(n_3) + 2Z_2(n_2, n_3)Z(n_1)Z(n_4) + 2Z_2(n_2, n_4)Z(n_1)Z(n_3) + \\ &2Z_2(n_3, n_4)Z(n_1)Z(n_2) + 4Z_2(n_1, n_2)Z_2(n_1, n_2) + 4Z_2(n_1, n_4)Z_2(n_2, n_3)\Big). \end{aligned} \tag{S119}$$

The results for larger $m$ proceed in a similar fashion.

At this point it is straightforward to include the penalty for multimerization, which (as discussed in Section S2) we set to $\Delta G_{\text{assoc}} = 4.09$ kcal/mol $- k_B T \log(\rho/1$ mol/L$)$. $Z_2$ is multiplied by one factor of $\exp(-\beta\Delta G_{\text{assoc}})$; $Z_3$ by two factors; $Z_4$ by three, etc.
